# Identification of an arabinopyranosyltransferase from *Physcomitrella patens* involved in the synthesis of the hemicellulose xyloglucan

**DOI:** 10.1101/241521

**Authors:** Lei Zhu, Murali Dama, Markus Pauly

**Affiliations:** Department of Plant and Microbial Biology, University of California, Berkeley, Berkeley, CA, 94720, USA; Institute of Plant Cell and Biotechnology, University of Dusseldorf, Dusseldorf, Germany

**Keywords:** Hemicellulose, Xyloglucan, glycosyltransferase, arabinopyranosyltransferase, *Physcomitrella patens*

## Abstract

The hemicellulose xyloglucan consists of a backbone of a β-1,4 glucan substituted with xylosyl moieties and many other, diverse sidechains that are important for its proper function. Many, but not all glycosyltransferases involved in the biosynthesis of xyloglucan have been identified. Here, we report the identification of an hitherto elusive xyloglucan:arabinopyranosyltransferase. This glycosyltransferase was isolated from the moss *Physcomitrella patens*, where it acts as a **<u>X</u>**yloglucan “**<u>D”</u>**-side-chain **<u>T</u>**ransferase (**XDT**). Heterologous expression of *XDT* in the *Arabidopsis thaliana* double mutant *mur3.1 xlt2*, where xyloglucan consists of a xylosylated glucan without further glycosyl substituents, results in the production of the arabinopyranose-containing “D” side chain as characterized by oligosaccharide mass profiling, glycosidic linkage analysis, and NMR analysis. In addition, expression of a related *Physcomitrella* glycosyltransferase hortholog of *XLT2* leads to the production of the galactose-containing “L” side chain. The presence of the “D” and “L” xyloglucan side chains in *PpXDT mur3.1 xlt2* and *PpXLT2 mur3.1 xlt2* transgenic plants, respectively, rescue the dwarfed phenotype of untransformed *mur3.1 xlt2* mutants to nearly wild-type height. Expression of *PpXDT* and *PpXLT2* in the Arabidopsis *mur3.1 xlt2* mutant also enhanced root growth.

## INTRODUCTION

The plant cell wall is a complex extracellular matrix composed of polysaccharides such as cellulose, hemicellulose, and various pectic polysaccharides, glycoproteins and the polyphenol lignin. The major hemicellulose xyloglucan (XyG) is found in all land plants and is especially abundant in the primary cell wall of dicots (1). XyG in the primary cell wall attaches to cellulose microfibrils non-covalently via H-bonds and its metabolism in the wall is thought to play a role in cell elongation (2, 3, 4). However, the precise molecular role of XyG in plant growth and development is not clear (5, 6, 7) as mutant plants lacking XyG do not exhibit an obvious growth phenotype (8). Initially it was thought that a particular XyG structure is plant species specific, but recently tissue specific structures within a plant species have emerged (9, 10, 11). XyG has not only been found in higher plants, but also in non-vascular plants such as liverworts and mosses (12).

XyG consists of a backbone of β-1,4 glucan substituted with xylosyl residues that are often further decorated with other sugar residues and/or acetyl-residues, leading to the discovery of more than 20 structurally different XyG side chains to date (13, 14, 15). Due to the structural diversity, a one-letter code has been established describing XyG side chains (16). According to this code G refers to an unsubstituted glucosyl backbone residue, while X depicts a xylosylated glucosyl residue as in α-D-xylose-6-β-D-glucose. X can be further extended on the xylosyl unit at *O-2* with galactosyl-, arabinopyranosyl-, galacturonsyl-, xylosyl- or arabinofuranosyl residues resulting in L, D, Y, U, and S side chains, respectively (12, 17, 18, 19, 20).

XyG assembly requires various glycosyltransferases (GTs) that add specific sugars to the extending polymer. Many GTs involved in XyG synthesis have been identified that belong to various Carbohydrate-Active Enzymes (CaZy) families based on gene-sequence homology (1, 21). One of the CaZy families involved in XyG sidechain biosynthesis is the GT47 family, including MUR3, XLT2, and XST (Fig. 1). MUR3 represents a XyG:galactosyltransferase, which adds a β-galactosyl-residue at the *O*-2 position to a xylosyl-residue resulting in the sidechain L (22). MUR3 transfers the galactosyl-moiety to a specific xylosyl-residue on the XyG chain leading to the occurrence of an XXLG oligosaccharide motive in XyG. In contrast, a related GT47 protein, XLT2, adds the galactosyl-residue to another xylosyl-residue leading to a XLXG motive indicating that these GTs exhibit regioselectivity (23). GT47 family members can also transfer galacturonic acid (XUT1) or arabinofuranosyl-moieties (XST) (17, 24) to the xylosyl-residue.

**Figure 1.**
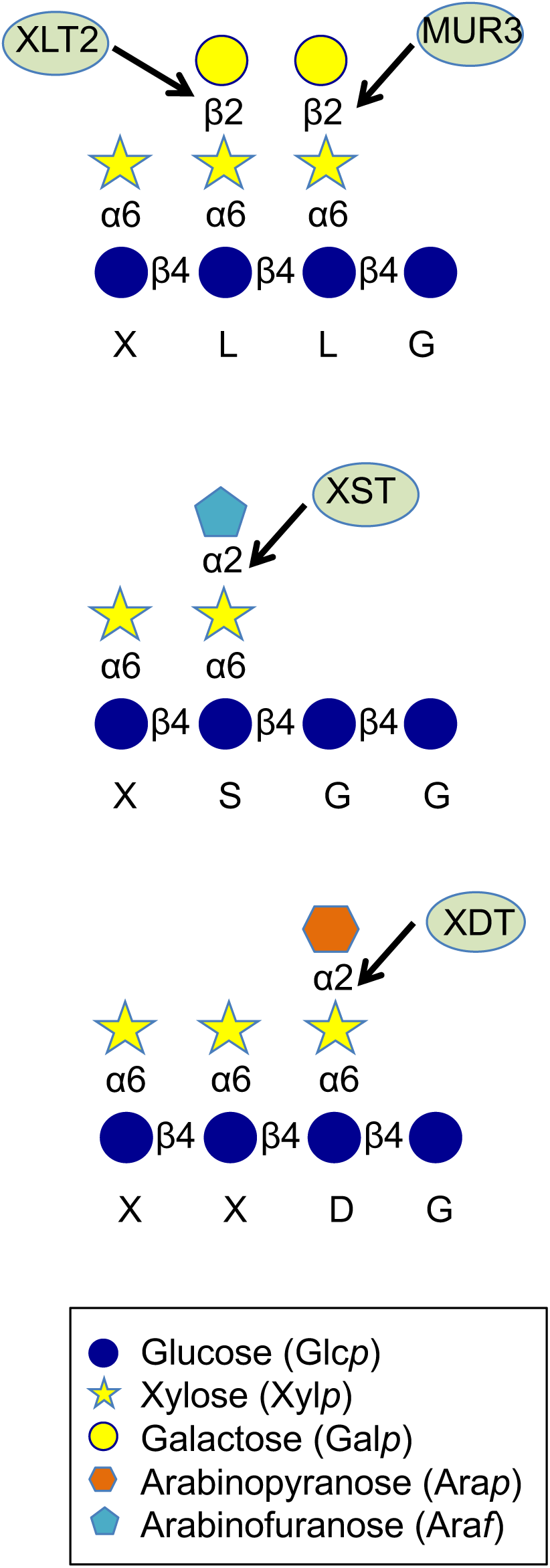
Xyloglucan oligosaccharide structures and GT47 glycosyltransferases involved in its synthesis. The xyloglucan one-letter code nomenclature is indicated below the structure. MUR3 and XLT2 – XyG:galactosyltransferases; XST – XyG:arabinofuranosyltransferase; XDT – XyG:arabinopyranosyltransferase

The moss *Physcomitrella patens* was found to contain XyG (12) with branched side chains containing galacturonosyl and arabinopyranosyl residues at the *O-2* position of their xylosyl residues (12). The arabinopyranosyl residue is unique as it has also been found in the XyG of lower plants such as the Lycophytes including Selaginella, Equisetales, Polypodiales and Cycadales (12, 18), but not in any gymnosperm or angiosperm plant to date.

To gain insights into the function of the XyG:arabinopyranosyl residue on XyG side chains, we describe here the identification of the responsible *Physcomitrella* arabinopyranosyltransferase present in the GT47 family. Because the simplest XyG side chain containing an arabinopyranosyl residue has been abbreviated with the one letter code D (1, 16), we named the responsible protein XDT (**X**yG **D** side chain **T**ransferase).

## RESULTS

### Identification of XyG-related GT47 Family Members in the Moss *Physcomitrella patens*

The amino acid sequence of the Arabidopsis XyG-related GT47 family member AtMUR3 was used as a bait to identify related GT candidates of *Physcomitrella* present in the Joint Genome Institute database Phytozome (phytozome.jgi.doe.gov). Based on amino acid sequence homology 13 *Physcomitrella* proteins were identified, which were also homologous to other, known GT47 XyG related genes from various species (AtXLT2, AtXUT, OsMUR3, OsXLT2, and SlXST; Fig. 2). Of the 13 Physcomitrella proteins, 6 members grouped closely in a MUR3 subclade. The other 7 Physcomitrella proteins fell into the XLT2 subclade that also included XST and XUT. Based on the location in the protein phylogenetic tree 9 non-redundant proteins were chosen for further, functional investigation (Pp1918, Pp42620, Pp201625, Pp2661, Pp21725, Pp173836, Pp156311, Pp110748, and Pp13057).

**Figure 2.**
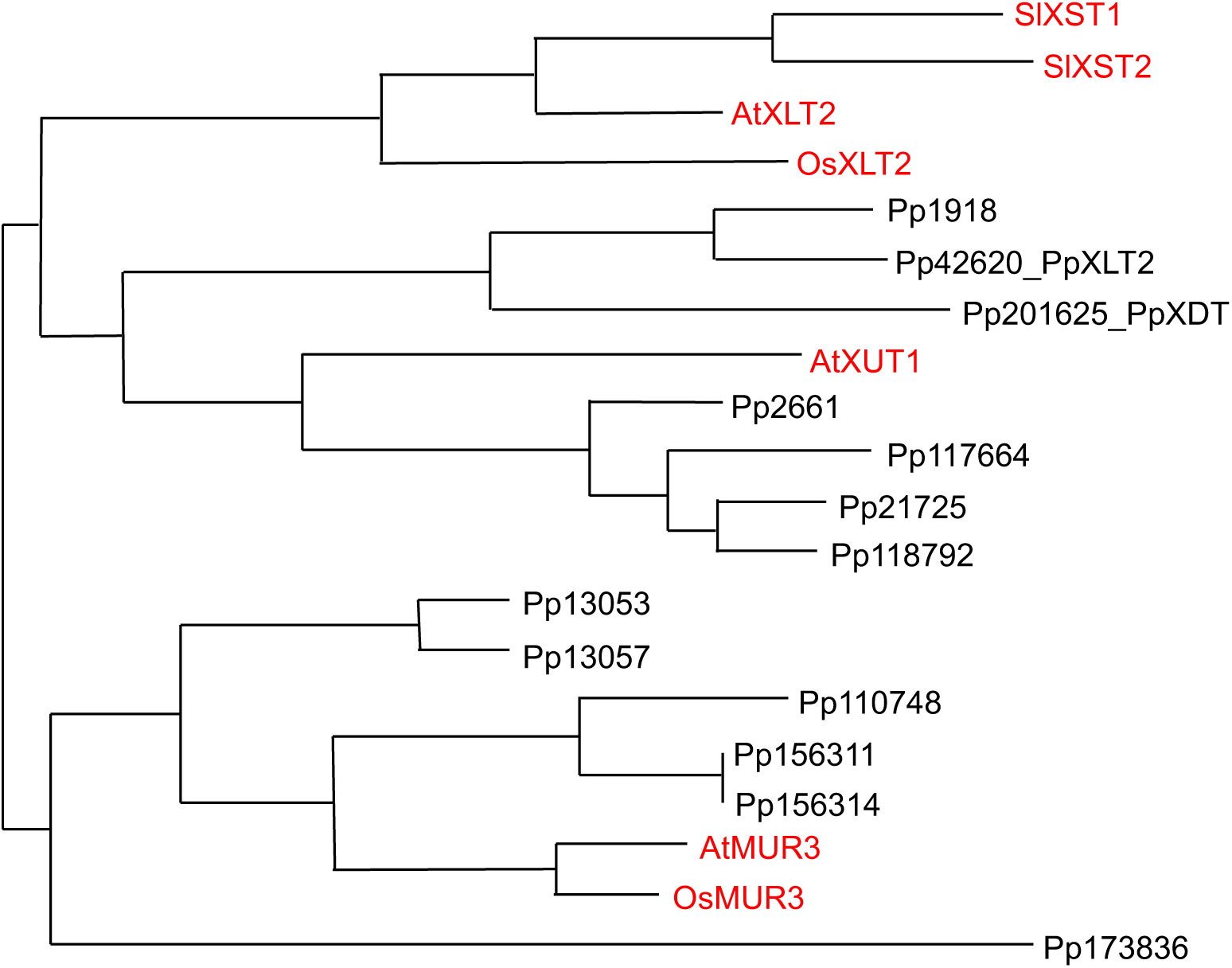
Phylogeny of XyG-related GT47 proteins. Red font – Protein sequences from known XyG:galactosyltransferases AtMUR3, OsMUR3, AtXLT2 and OsXLT2, the galacturonosyltransferase AtXUT, and the arabinofuranosyltransferases SlXSTl and SlXST2. Black font – *Physcomitrella patens* proteins obtained from a BlastP search against Phytozome database. Phylogenetic tree built by PhyML.

### Functional Complementation in Arabidopsis and Characterization of XDT

To assign GT functions to the 9 selected Physcomitrella GT47 family members heterologous expression of individual proteins in the Arabidopsis double mutant *mur3.1 xlt2* was pursued. XyG derived from the various complemented Arabidopsis plants was analyzed by oligosaccharide mass profiling (OLIMP) (25), whereby XyG was solubilized from wall materials using a xyloglucan specific endoglucanase and the resulting XyG oligosaccharide mixture was analyzed by MALDI-TOF mass spectrometry (Fig. 3). The OLIMP profile of untransformed *mur3.1 xlt2* mutant plants shows the occurrence of a single oligosaccharide motive with a m/z of 1,085 representing the XyG oligosaccharide XXXG consisting only of the glucan backbone with xylosyl moieties but no further substitutions. This OLIMP profile was retained when seven of the Physcomitrella genes were constitutively expressed in Arabidopsis *mur3.1 xlt2* indicating that in Arabidopsis these genes are not involved in XyG biosynthesis.

**Figure 3.**
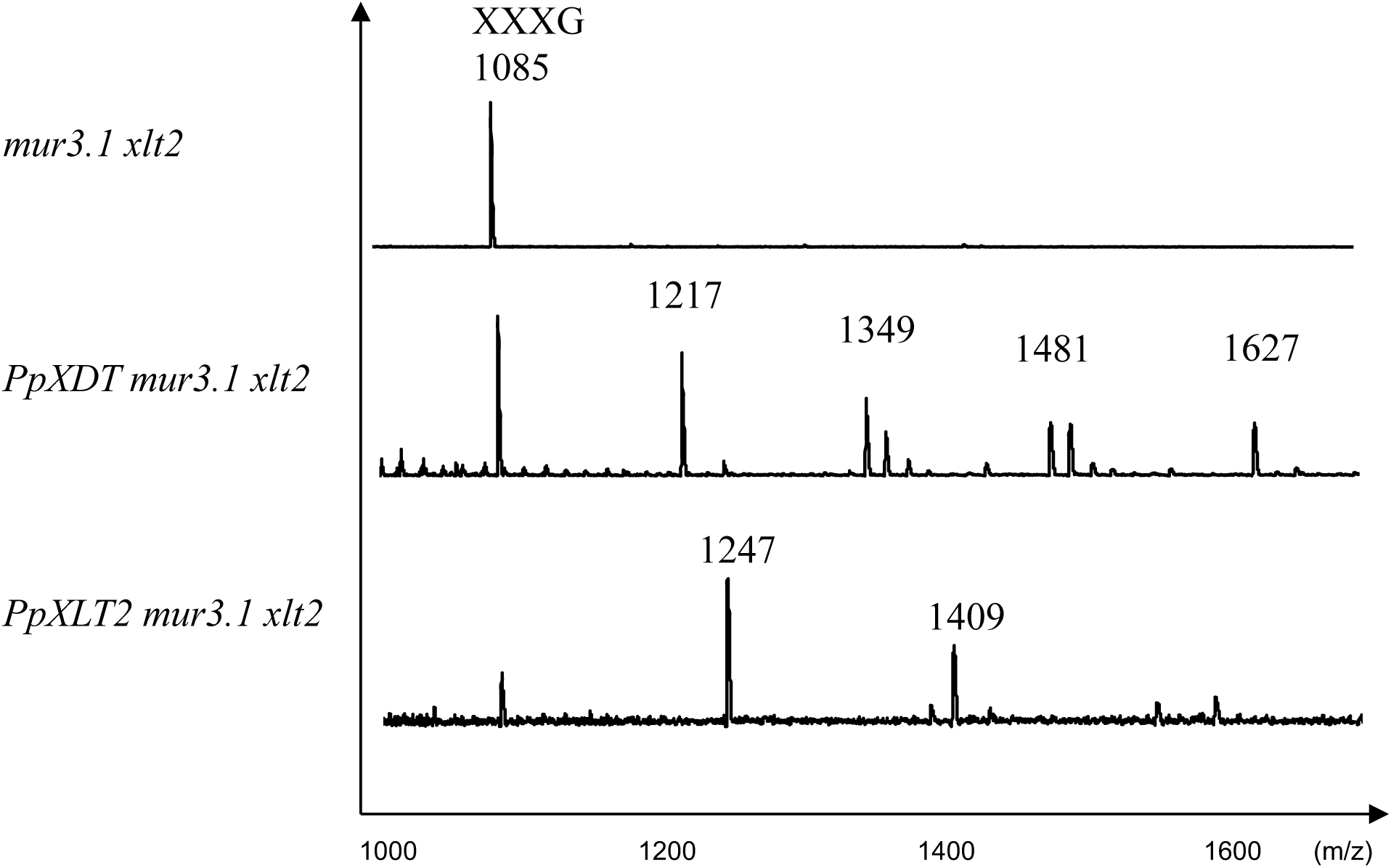
XyG oligosaccharide mass profiling by MALDI-TOF MS. XyG oligosaccharides derived from leaf tissue of the Arabidopsis double mutant *mur3.1 xlt2*, transgenic lines expressing *PpXDT* or *PpXLT2* in *mur3.1 xlt2*. Numbers indicate m/z and their potential structure is shown in supplemental table 1. m/z 1085 represents the known XyG oligosaccharide structure XXXG.

However, expression of *Pp201625 (PpXDT)* in *mur3.1 xlt2* resulted in a XyG that contained an oligosaccharide of m/z 1,217 indicating that this GT affects XyG biosynthesis and is responsible for adding an additional pentosyl residue to XXXG in *mur3.1 xlt2* (Fig. 3). Moreover, 5 additional XyG oligosaccharides were observed when *PpXDT* was expressed in the double mutant *mur3.1 xlt2* (Fig. S1). These ions with an m/z of 1,349, 1,363, 1,481, 1,495, and 1,627 correspond to oligosaccharide structures consisting of 4 hexoses and 5 pentoses (H4P5), H4P4 with an additional deoxysugar, likely to be fucose, H4P6, H4P5 with an additional fucose, and H4P6 with a fucose, respectively. Mass spectrometry neither gives an indication which kind of pentose was added nor where the pentose would be attached. To determine the fine structure of the dominant novel XyG oligosaccharide (m/z 1,217) the oligosaccharide was isolated/ enriched by subjecting the XyG oligosaccharide mixture obtained from wall material of *Pp201625 mur3.1 xlt2* to High Performance Anion Exchange Chromatography with Pulsed Amperometric Detection (HPAEC-PAD; Fig. 4). Oligosaccharide(s) with a molecular mass of m/z 1,217 eluted at ~13.2 min and were collected for further analysis. Some impurities of the XyG oligosaccharide XXXG were present in the collected fraction due to its adjacent elution. The retention time of the novel oligosaccharide was found to be the same as a well characterized XyG oligosaccharide isolated from *Selaginella kraussiana* (18), termed XXDG, a XyG oligosaccharide containing an arabinopyranosyl residue (Fig. 4). The isolated/enriched oligosaccharide with a m/z of 1217 isolated from *Pp201625 mur3.1 xlt2* was subjected to glycosidic linkage analysis (Table 1). The results indicate the presence of t-Arap and 2-Xylp in an equal ratio supporting the hypothesis that Arap is attached to *Xylp* at *O-2* thus representing the structure of XyG D side chain. No t-Ara*f* was detected. To gain further insights into this structure ^1^H NMR was performed (Fig. 5, Additional Fig. S1). Based on previously observed chemical resonances (12) the data confirmed the presence of an anomeric signal of an α-linked arabinopyranosyl residue (chemical shift of 4.488 ppm). In addition, the corresponding substituted 2-α-Xylp residue was identified with a chemical shift of 5.133 ppm. The observed chemical shifts are therefore consistent with the presence of the oligosaccharide XXDG (Fig. 5). Hence, Pp201625, when expressed in the Arabidopsis mutant *mur3.1 xlt2*, plays a role in transferring arabinopyranose to XXXG generating a XyG D side chain and was thus termed XyG D-side-chain Transferase (XDT).

**Table 1.** Glycosidic linkage analysis of XyG oligosaccharide fraction with a m/z 1217 (with contamination of XXXG) from leaf walls of *PpXDT mur3.1 xlt2*.

| Sugar moiety | Abundance (%) |
| --- | --- |
| t-Arap | 2.88 |
| t-Xyl | 38.20 |
| 2-Xyl | 2.88 |
| 6-Glc | 14.57 |
| 4-Glc | 13.29 |
| 4,6-Glc | 28.18 |

Note: t-Arap, terminal arabinopyranose; t-Xylp, terminal xylopyranose; 2-Xyl, 2-linked xylose; 6-Glc, 6-linked glucose; 4-Glc, 4-linked glucose; 4,6-Glc, 4,6-linked glucose

**Figure 4.**
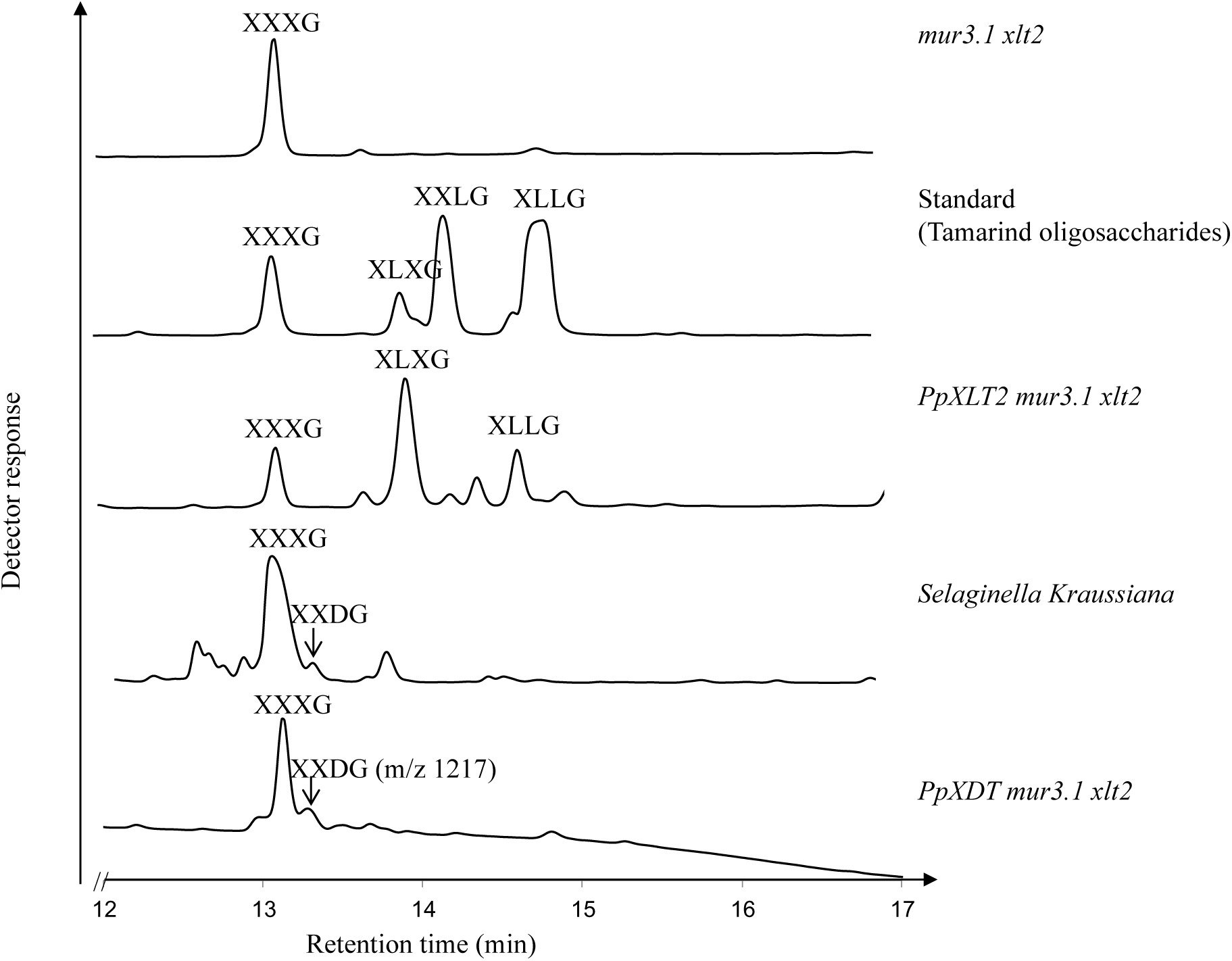
XyG oligosaccharide separation by HPAEC-PAD. Peaks were assigned based on retention times from published work (Schultink et al., 2013; Hsieh & Harris, 2012; Megazyme Inc., Ireland) as well as assignment based on mass spectrometry. The fractions containing m/z 1217 (XXDG) was collected and further analyzed by glycosidic linkage analysis (Table 1) and NMR spectroscopy (Fig. 5).

**Figure 5.**
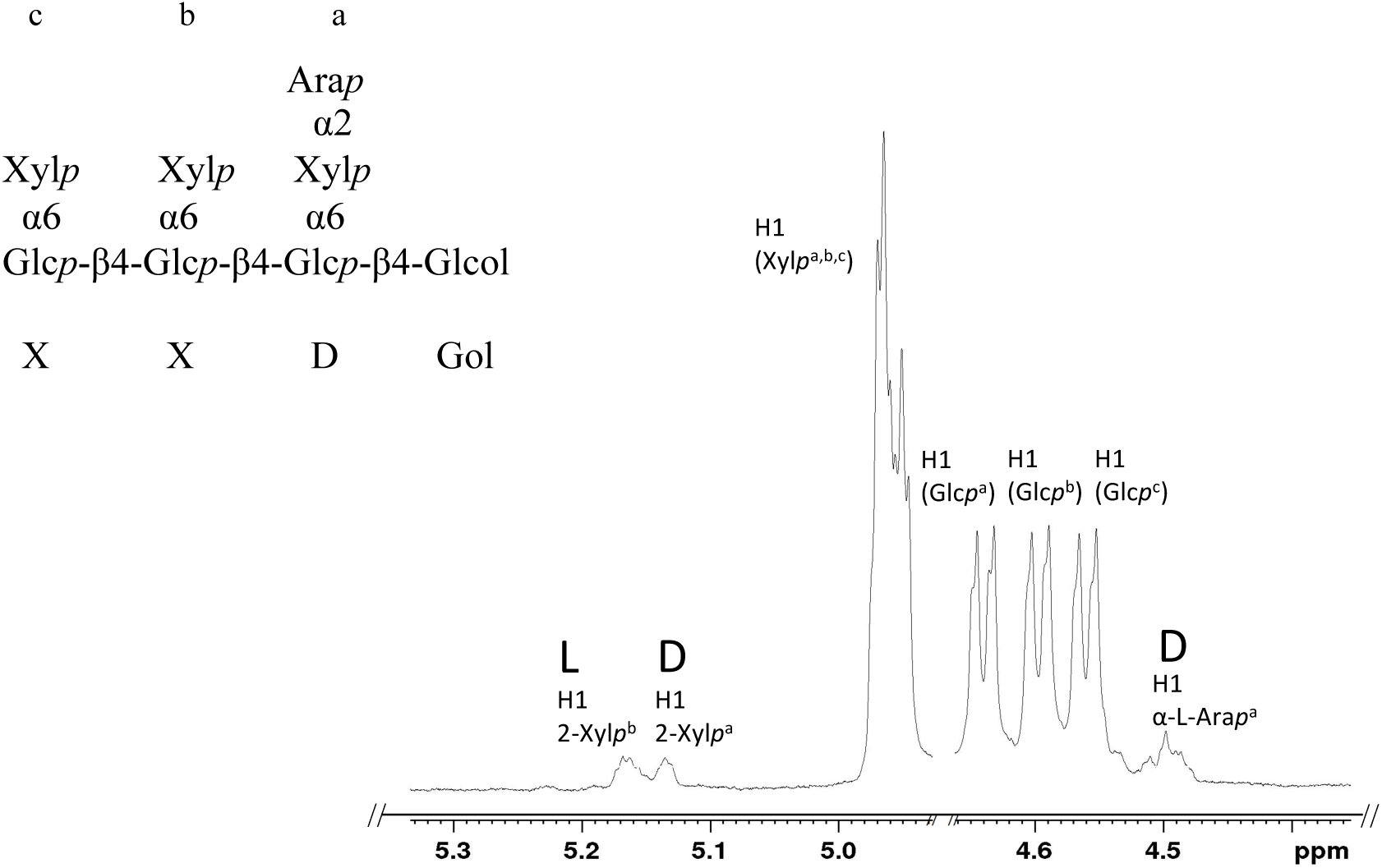
Anomeric region of the ^1^H NMR spectra of the reduced XyG oligosaccharide XXDGol contaminated with XXXGol. ^1^H NMR peaks were labelled according to CCRC (ccrc.uga.edu) database and the literature (Pena et al., 2008). The putative structure of the oligosaccharide is shown in the upper left corner indicating the order of the side-chains on top and the one-letter code below.

### Heterologous expression of *PpXLT2*

The Physcomitrella GT47 family also contains Pp42620, a protein that phylogenetically belongs to the AtXLT2 subclade (Fig.2). Expression of Pp42620 in Arabidopsis *mur3.1 xlt2* resulted in the generation of various XyG oligosaccharides, when the transgenic plants were analyzed by oligosaccharide mass profiling. The oligosaccharides with a m/z of 1,247 represents XXXG plus an additional hexose – the minor new oligosaccharide with a m/z of 1,409 represents XXXG plus 2 hexoses (Fig. 3). To determine the fine structure of the dominant XyG oligosaccharide (m/z 1,247) the XyG oligosaccharide mixture generated from wall materials of *Pp42620 mur3.1 xlt2* was analyzed by HPAEC-PAD. Compared to previously published data (26) and using commercially available tamarind XyG oligosaccharides as standards the novel oligosaccharide exhibited the same retention time as tamarind XyG oligosaccharide XLXG (Fig. 4). Also oligosaccharides with the retention time of XXLG and the double galactosylated XLLG oligosaccharide were present suggesting that *Pp42620*, when expressed in Arabidopsis, exhibits mainly XLT2 activity, hence its name PpXLT2, but also to some extent MUR3 galactosyltransferase activity.

### Growth Phenotypes of *PpXDT mur3.1 xlt2* and *PpXLT2 mur3.1 xlt2*

The structure of XyG has been shown to affect vegetative growth. For example, the Arabidopsis double mutant *mur3.1 xlt2* containing XyG entirely composed of XXXG units exhibits dwarfism (9, 24) (Fig. 6). When *PpXDT* and *PpXLT2* were expressed in *mur3.1 xlt2* using a constitutive promotor vegetative (stem) growth was restored to nearly normal heights in most of the lines (Fig. 6 A, B). The Arabidopsis double mutant *mur3.1 xlt2* exhibits also shorter roots (17) (Fig. 7 A, B). Expression of PpXDT and *PpXLT2* in the double mutant leads to root growth that is not significantly different than Arabidopsis WT.

**Figure 6.**
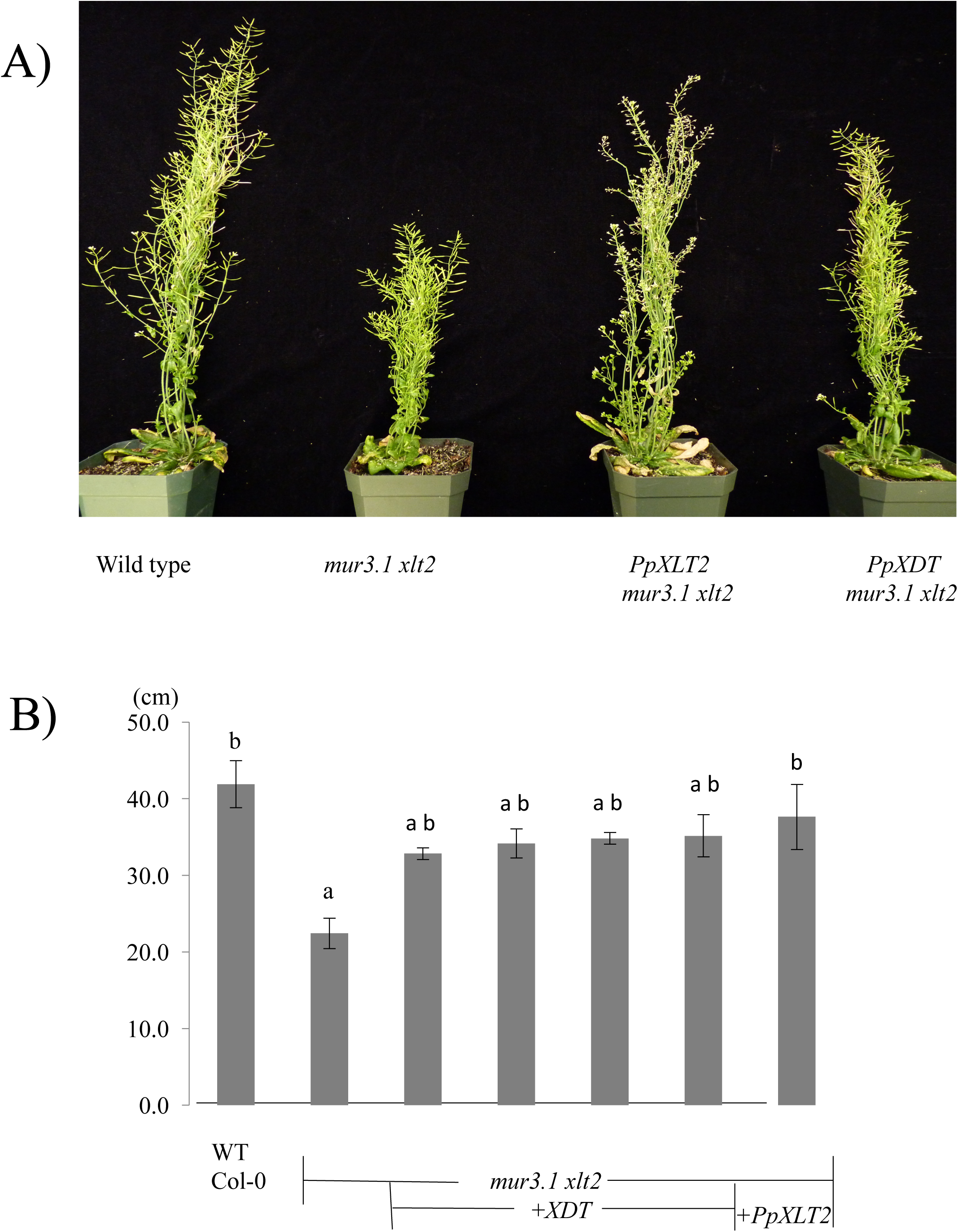
A) Growth habit of 8-week old Arabidopsis plants. B) Height of Inflorescence stems of 8-week old Arabidopsis plants. n≥3. Small letters – ANOVA analysis.

**Figure 7.**
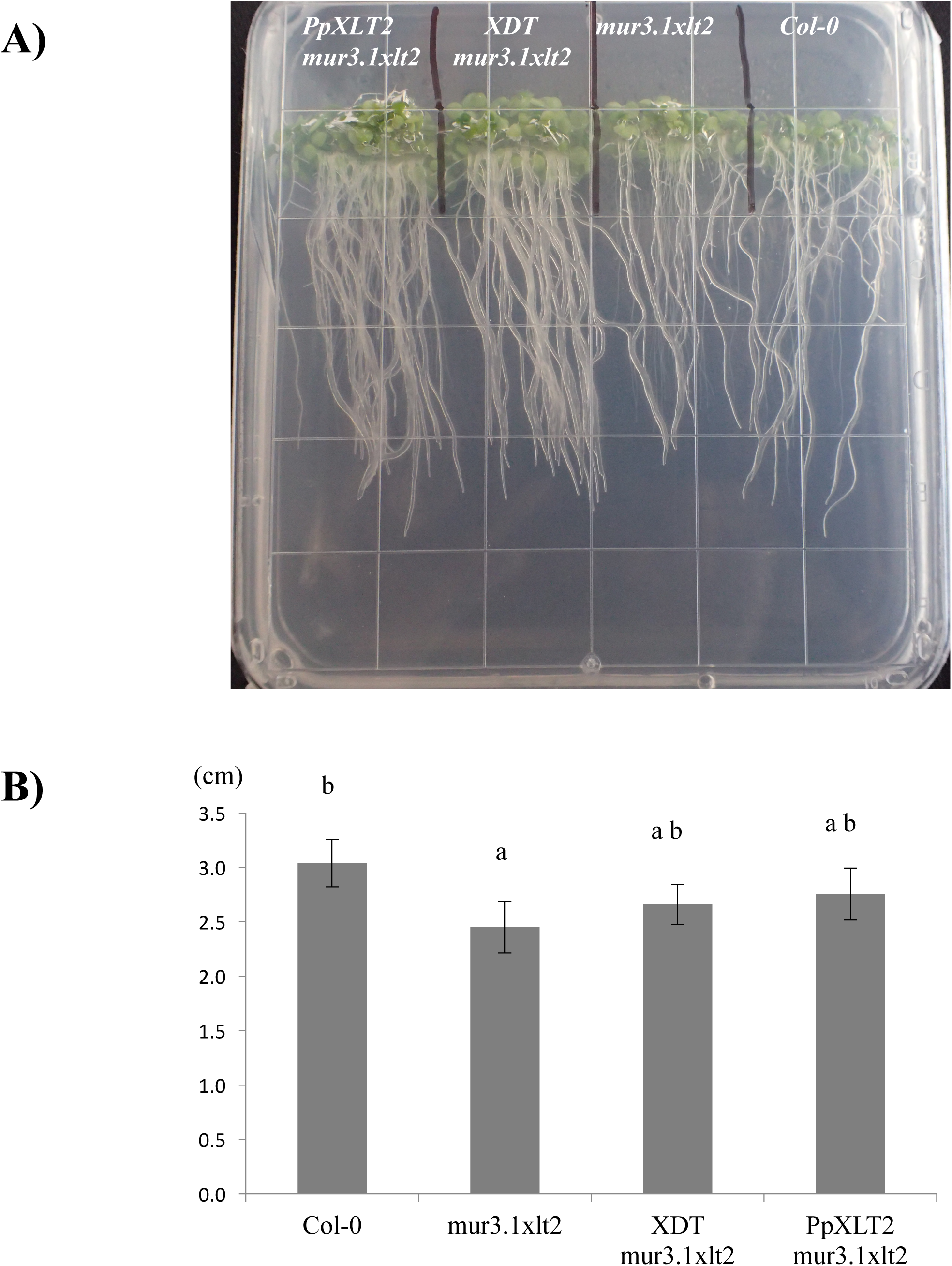
Root growth of Arabidopsis plants. A) Root habit B) Root length of 8-day-old seedlings, n=10, small letters – ANOVA analysis.

## DISCUSSION

### Identification of a XyG:Arabinopyranosyltransferase (XDT)

The xylosyl residue of XyG is often substituted at the *O-2* position with a variety of glycosyl residues including galactosyl, galacturonosyl, xylopyranosyl, arabinofuranosyl or arabinopyranosyl moieties (1). Using *in vitro* assays, loss-of-function mutants and functional complementation approaches in Arabidopsis have led to the successful identification of many of the responsible GTs including two galactosyltransferases MUR3 and XLT2 (22, 23), a galacturonsyltransferase XUT1 (17), an arabinofuranosyltransferase XST (24) and as identified and characterized in this study an arabinopyranosyltransferase XDT.

The availability of the *Physcomitrella patens* genome (27) allowed us to identify this XyG:arabinopyranosyltransferase XDT. Expression of the *XDT* gene from Physcomitrella led to synthesis of the XyG D side chain in the *Arabidopsis* double mutant *mur3.1 xlt2* as evidenced by XyG analysis by MALDI-TOF MS, HPAEC-PAD, linkage analysis and NMR. When a MUR3 ortholog from rice is expressed in the Arabidopsis *mur3.1 xlt2* double mutant XyG does not only become galactosylated, it also becomes fucosylated (9) as the galactosylated side chain L is the required acceptor substrate for the XyG:fucosyltransferase (28) resulting in the F sidechain. Here, expression of *XDT* resulted not only in arabinosylated side chains, but also to a much lesser extent in side chains containing an additional deoxyhexose. This data is consistent with the occurrence of a fucosylated D side chain termed E, which has been observed in XyG derived from *Equisetum hyemale* and *Selaginella kraussiana* (12, 18). The Arabidopsis XyG:fucosyltransferase AtFUT1/AtMUR2 is apparently not only able to transfer fucosyl residues to galactosyl but also arabinopyranosyl residues. Similar to previous reports (12, 18) acetylated versions of the D side chain were not observed, indicating that the Arabidopsis XyG:O-acetyltransferase AtAXY4/AtAXY4L (29) specifically adds acetyl substituents to galactosyl residues.

The galactosyltransferases AtMUR3 and AtXLT2 act regiospecific, i.e. they transfer the galactosyl moiety to a specific xylosyl residue leading to the generation of XXLG or XLXG, respectively. The expression of *PpXDT* in Arabidopsis also lead to XyG oligosaccharides that in addition to arabinopyranosyl residue contain additional pentoses. Although the nature and position of these additional pentosyl residues remain to be determined it seems clear that XDT is more promiscuous in nature than MUR3/XLT2. Either the enzyme can transfer other pentoses than arabinopyranoses such as arabinofuranoses or more likely it can add arabinopyranoses to different positions on XyG resulting not only in the XyG oligosaccharide XXDG (m/z 1,217, Table S1) but also XDXG (m/z 1,217), XDDG (m/z 1,349), and even DDDG (m/z 1,481) and their fucosylated versions XXEG (m/z 1,363), XDEG (m/z 1,495), and DDEG (m/z 1,627). Ions of all these oligosaccharides were present when *PpXDT* was expressed in the *mur3.1 xlt2* mutant (Table S1). Thus, XDT does not seem to act regiospecific.

### Functional Conservation of XyG-related Genes in the GT47 Family

The GT47 family is a large carbohydrate-active enzyme (CaZy) family involved in cell wall biogenesis containing a subclade represented by the XyG:galactosyltransferase AtMUR3 (22) and includes XLT2, another XyG:galactosyltransferase (23). The identified Physcomitrella XyG:arabinopyranosyltransferase XDT also belongs to this subclade, as it forms the same glycosidic linkage on the same acceptor substrate albeit utilizing a different donor substrate (UDP-L-arabinopyranose). Taken together with the characterized functions of other GT47 members such as XUT1 from *Arabidopsis* and XST from tomato, members of the GT47 MUR3 subclade in land plants have evolved in transferring a glycosyl moiety to the xylosyl residue of XyG at *O*-2.

Within this clade another functional XyG GT was identified in *Physcomitrella patens*, PpXLT2. Analysis of the XyG present in the constitutive expression of *PpXLT2* in *Arabidopsis mur3.1 xlt2* resulted in the occurrence of XLXG indicating that PpXLT2 in *Arabidopsis* can carry out the same function as AtXLT2. However, in addition XXLG was formed including its fucosylated version XXFG. Thus, unlike AtXLT2 from Arabidopsis (23), SlXLT2 from tomato (24), and OsXLT2 from rice (9) the Physcomitrella ortholog PpXLT2 does not act regiospecific, it also exhibits MUR3 activity. Therefore, while the function of XLT2 is functionally conserved across land plants including bryophytes, its regioselectivity of this GT has apparently evolved later as it has to date only been observed in angiosperm species.

### XyG side chains impact aerial and root growth

Arabidopsis mutant *mur3.1* contains a point mutation in the *MUR3* gene and renders it inactive (22). As a result, XyG in this mutant does not contain the XXLG oligosaccharide, but still the galactosylated XLXG motive. Mutant plants show normal plant growth except for minor effects in trichome morphology (22). However, the double mutant *mur3.1 xlt2*, whose XyG does neither contain XXLG nor XLXG oligosaccharides, but consists entirely of XXXG oligosaccharides displays a dwarfed phenotype (30). Complementing this mutant with various XyG GT47 genes results in the rescue of the growth phenotype. This rescue has been observed with the expression of *MUR3* and *XLT2* from a variety of species such as from rice *OsXLT2* (9), tomato *SIXLT2* (24), or as shown here from Physcomitrella *PpXLT2*. Moreover, arabinofuranosylation by expressing XST (24) and as shown here arabinopyranosylation through XDT also restores the phenotype of the double mutant, not only the growth of vegetative tissue, but also root growth. This indicates that galactosylation or the occurrence of the L side chain is not required for normal growth, but that alternative substitutions such as arabinofuranosylation and arabinopyranosylation resulting in the S and D side chains, respectively, suffice for normal plant growth. It is known that XyG that consists only of XXXG self-aggregates and precipitates *in vitro* (31, 32). Such precipitation of non-galactosylated XyG in the *mur3.1 xlt2* mutant might occur already during its biosynthesis in the Golgi-apparatus of impacting the endomembrane system function. Indeed, the Arabidopsis *mur3.3* mutant, an insertional knockout mutant, exhibits severe dwarfism with a concomitant aggregation of endomembranes and intracellular accumulation of polymers (30). However, when the *mur3.3* mutant is crossed with the XyG-lacking *xxt1 xxt2* mutant the resulting mutant plant exhibits not only again a normal growth phenotype but also a normal endomembrane morphology. XyG is still lacking in these plants. Hence a structurally abberant XyG with low or lacking galactosylation is detrimental to plant development, whereas a lack of XyG is not.

## METHODS

### Plant growth

Seeds of Arabidopsis wild-type Col-0, *mur3.1 xlt2* (24) and the transgenic plants generated here were germinated either in soil pots or on half MS agar plates. Plants were grown in a Percival growth chamber at 21°C under 16/8 hour light/dark cycle with 70% humidity.

### Phylogenetic analysis

*Pyscomitrella* XyG-related GTs were identified by using AtMUR3 as a template for a BlastP search of the *Physcomitrella* phytozome database (version 10.1). Alignment of the XyG-related GT47 proteins was achieved by MUSCLE alignment and construction of a phylogeny tree using PhyML (33, 34).

### Gene constructs and plant transformation

Physcomitrella candidate genes were amplified either by PCR from genomic DNA or by RT-PCR from total RNA extracted from 1-week-old protonemal tissue and 1-month-old gametophytes of *Physcomitrella patents*. Primer sequences used for cloning are listed in the additional table S2. The amplified genes were cloned into the expression vector pORE-E4, which was transformed to *Agrobacterium tumefaciens* strain GV3101, and subsequently transformed to Arabidopsis via the floral dip method (35). Three generations of transgenic *PpXDT* and *PpXLT2* plants were selected on half MS agar (0.8%) plates containing 60pμg ml^-1^ Kanamycin. Germinated seedlings were then move into soil for continuous growth.

### Analysis of xyloglucan

XyG oligosaccharides were extracted from leaf tissue of Arabidopsis Col-0, *Atmur3.1 xlt2, PpXDT mur3.1 xlt2, PpXLT2 mur3.1 xlt2*, and *Selaginella Kraussiana* by alcohol insoluble residue (AIR) preparation followed by xyloglucanase digestion (36) and subsequent XyG oligosaccharide profiling by MALDI-TOF MS and HPAEC-PAD as described (23, 24).

### Purification of xyloglucan oligosaccharide XXDG

Extraction of the XyG oligosaccharide XXDG (m/z 1217) from transgenic plants was performed according to methods described in Schultink et al., 2013. However, the reduction of the oligosaccharides by sodium borohydride was performed after separation of the oligosaccharides by HPAEC-PAD. The reduced oligosaccharides were neutralized, de-salted using a ENVI-CARB reverse phase column (Sigma Aldrich, USA) and freeze dried in a lyophilizer.

### Glycosyl linkage analysis

Glycosidic linkage analysis of XyG oligosaccharides was performed as described (24).

### NMR analysis

Reduced oligosaccharides were dissolved in 0.3 ml of D_2_O (99.9%, company?), freeze dried and dissolved again in 0.3 ml of D_2_O (99.9%) containing 0.05 % of 3-(trimethylsilyl)-propionic-2,2,3,3-d4 acid sodium salt). The ^1^H NMR spectra were recorded on a Bruker Avance 600 MHz NMR spectrometer equipped with an inverse gradient TXI ^1^H/^13^C/^15^N Cryoprobe at 298 K. All chemical shifts were referenced relative to 3-(trimethylsilyl) propionic-2,2,3,3-d4 acid (0.00 ppm for ^1^H). The NMR data processing and analysis was performed using Bruker’s Topspin 3.1 software.

## Abbreviations

XyG: Xyloglucan
XEG: Xyloglucan-specific endoglucanase
GT: Glycosyltransferase
CaZy: Carbohydrate-active enzyme
XDT: Xyloglucan D-side-chain Transferase
OLIMP: Oligosaccharide mass profiling
MALDI-TOF: Matrix-Assisted Laser Desorption/ Ionization Time of Flight Mass Spectrometry
HPAEC-PAD: high-performance anion-exchange chromatography withpulsed amperometric detection
GC-MS: Gas chromatography mass spectrometer
AIR: Alcohol insoluble residue
t-Ara*p*: terminal arabinopyranose
2-Xyl*p*: 2-xylopyranose
t-Ara*f*: terminal arabinofuranose

## Supplemental Figures

**Table S1.**
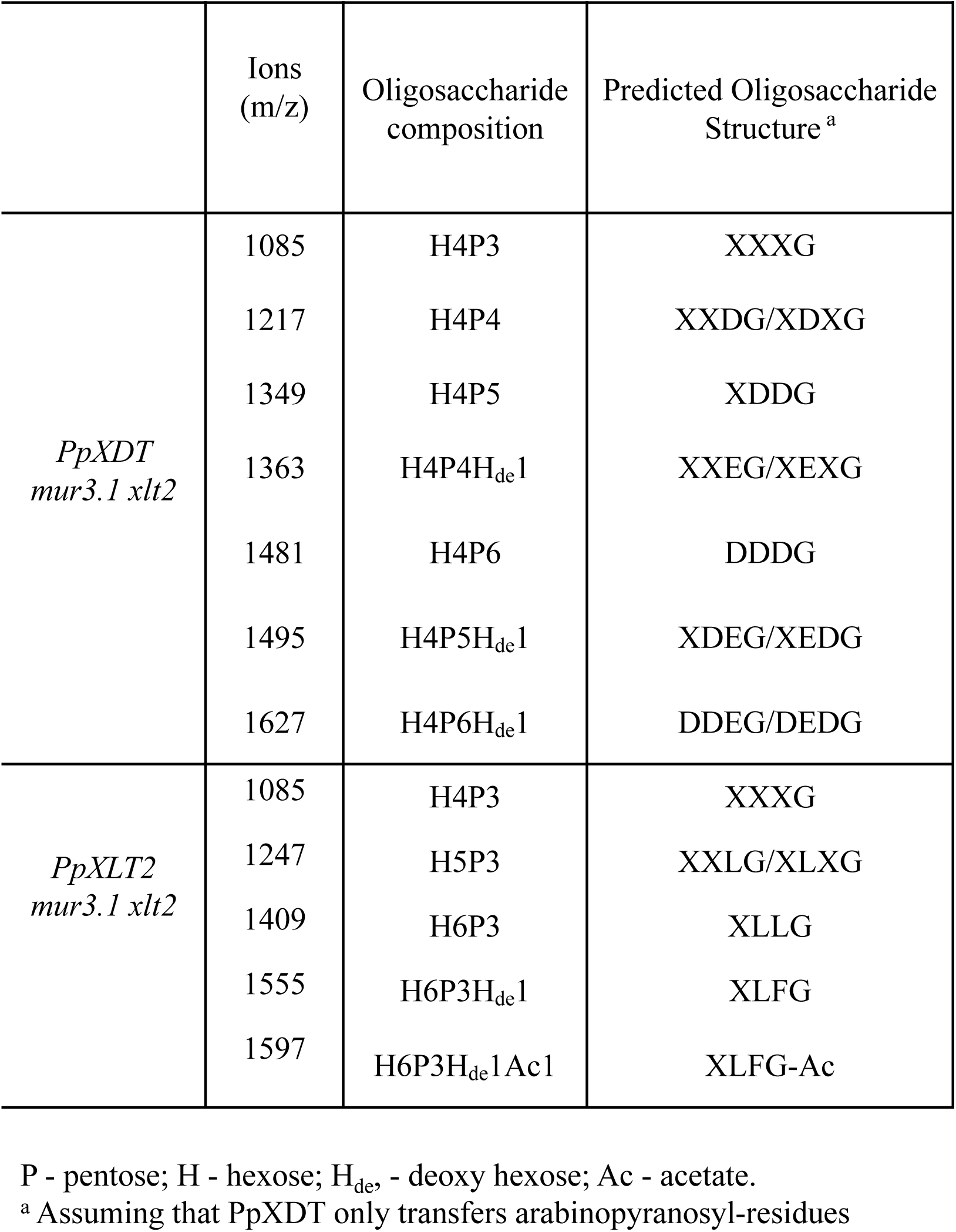
The XyG oligosaccharide composition and relative abundance of ions (m/z) seen in Fig. 3.

**Table S2.**
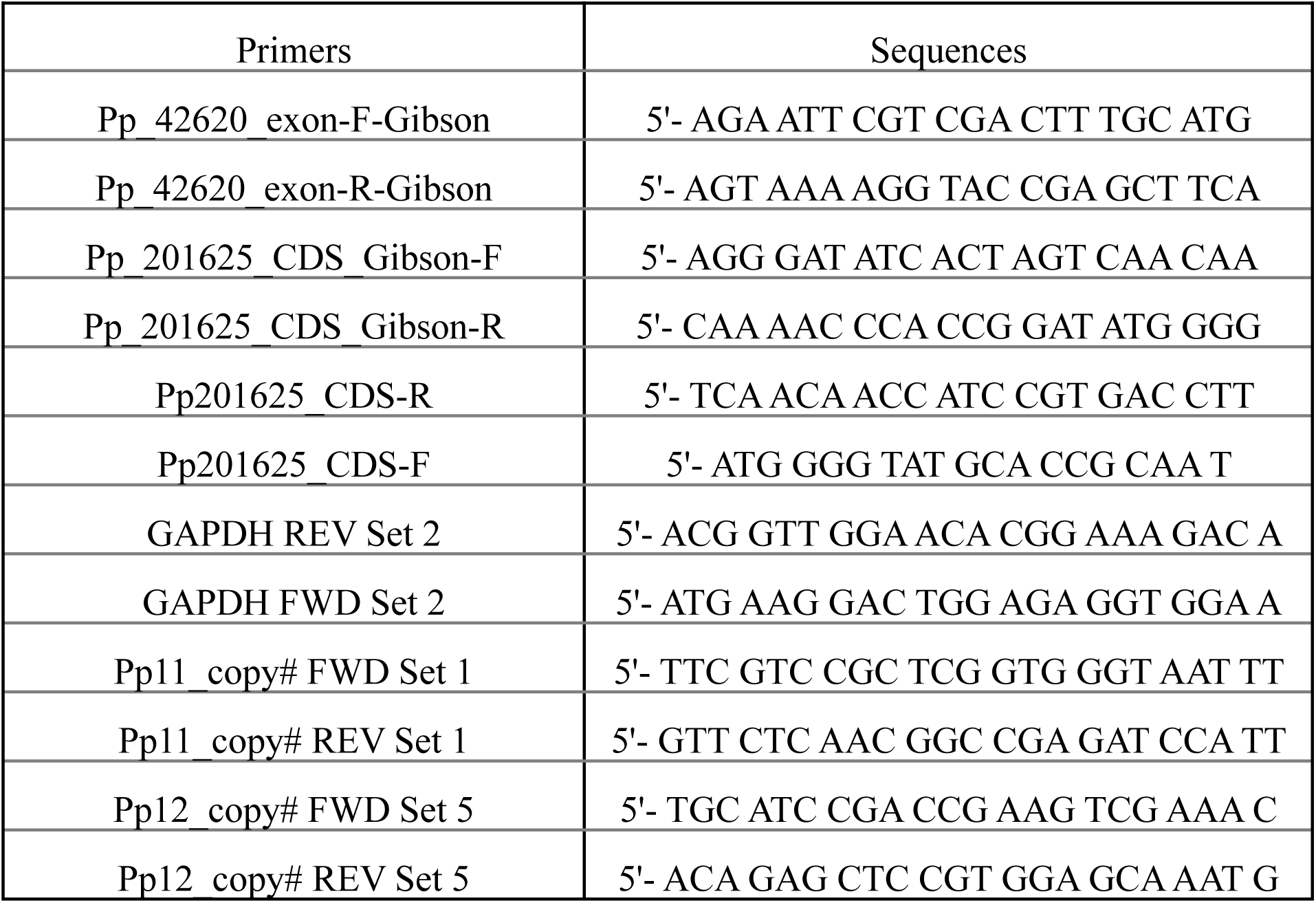
Primers for PCR amplification of genes and quantitative RT-PCR.

**Figure S1: Figure S1.**
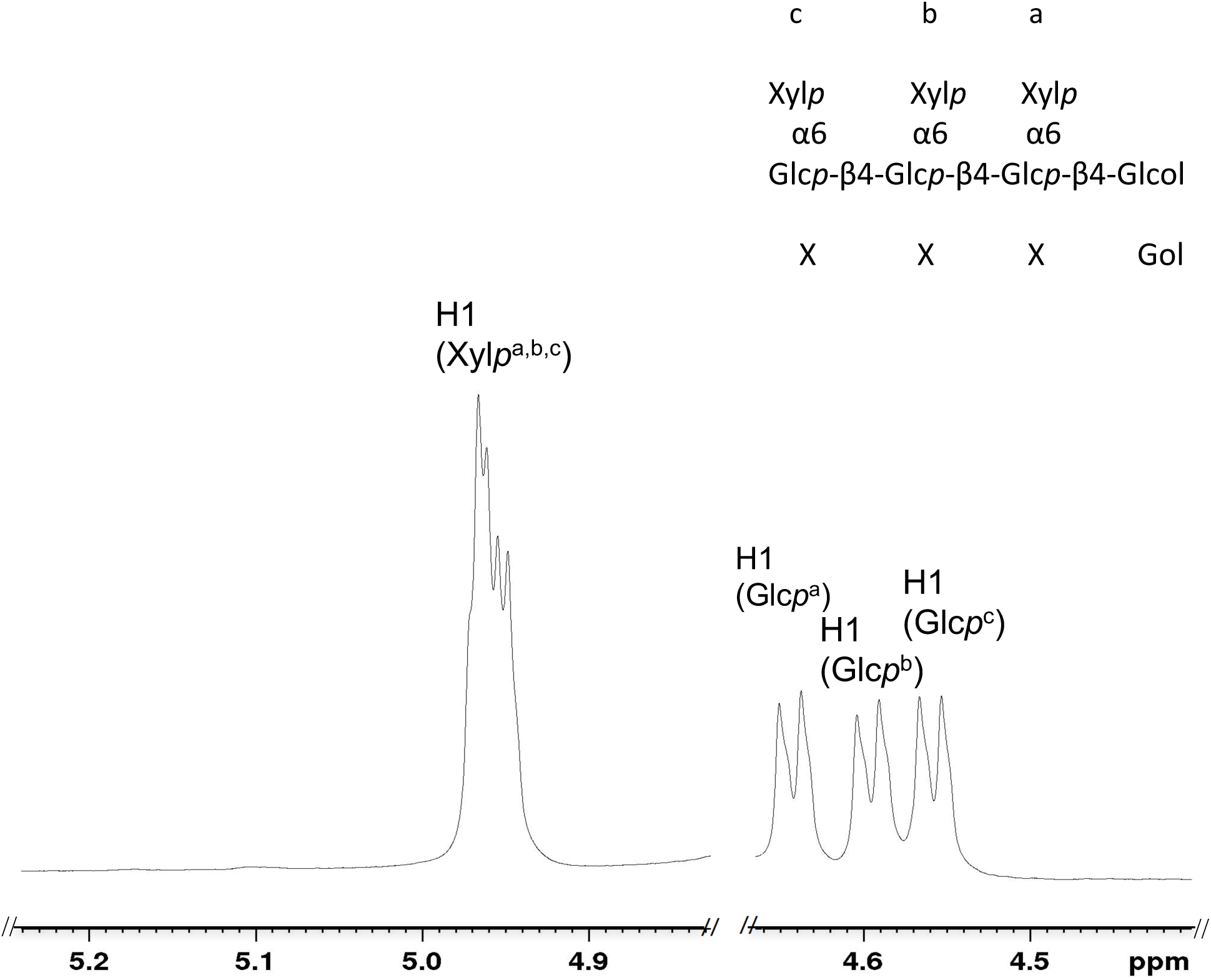
Anomeric region of the ^1^H NMR spectra of the oligosaccharide XXXG obtained from *mur3.1 xlt2*. ^1^H NMR peaks were labelled according to CCRC (ccrc.uga.edu) database. The putative structure of the oligosaccharide is shown in the upper right corner indicating the order of the side-chains on top and the one-letter code below.

## Declarations

### Ethics approval and consent

The leaf samples used in this study were collected from growth chamber in our lab and from botanical garden at University of California, Berkeley with permission of the curator. The experimental research was undertaken in accordance with local guidelines. For access to the plants, please contact the corresponding author.

### Consent for publication

Not applicable

### Availability of data and materials

Constructs described in this work and datasets analysed during the current study are available from the corresponding author upon request.

## Acknowledgements

We thank Dr. Tom Kleist and Professor Sheng Luan for providing moss protonemal and gametophyte tissue of Physcomitrella, and the botanical garden of the University of California, Berkeley for *Selaginella kraussiana*.

## Authors Contributions

LZ performed the experiments, analyzed the results and wrote the manuscript, MD performed the NMR experiments and wrote the manuscript, MP conceived the work, analyzed the data, and wrote the manuscript.

